# BIOPOINT: A particle-based model for probing nuclear mechanics and cell-ECM interactions via experimentally derived parameters

**DOI:** 10.1101/2025.04.22.650069

**Authors:** Sandipan Chattaraj, Julius Zimmermann, Francesco Silvio Pasqualini

**Affiliations:** Synthetic Physiology Lab, Department of Civil Engineering and Architecture, University of Pavia, 27100, Pavia, Italy

**Keywords:** particle-based model, deformable nucleus, cell-ECM interactions, nuclear mechanics, single cell indentation

## Abstract

Morphogenesis involves biochemical and biomechanical interactions across multiple spatial and temporal scales. Experimental studies alone struggle to resolve these dynamics, necessitating computational models. Among these, subcellular element modeling (SEM) has proven helpful in simulating cellular and tissue-scale emergent behaviors. However, traditional SEM frameworks lack explicit representations of nuclear mechanics and cell-extracellular matrix (ECM) interactions, limiting their ability to capture key biology.

Here, we introduce **BIOPOINT**, a particle-based computational framework that extends SEM by incorporating (1) a deformable, multi-particle nucleus to simulate nuclear stress and strain distributions and (2) an explicit ECM representation using a structured array of static particles interacting via tunable adhesive potentials. To ensure biological relevance, we calibrated BIOPOINT’s parameters against single-cell indentation experiments, overcoming prior limitations of ad hoc parameter selection in SEM. We validate BIOPOINT by comparing simulated cell behaviors to experimental observations in three key scenarios: (i) single-cell indentation, demonstrating agreement with force-time curves from atomic force microscopy (AFM) studies; (ii) cell spreading on ECM micropatterns, confirming that nuclear deformation follows ECM constraints; and (iii) nuclear deformation during confined migration, showing BIOPOINT predicts nuclear shape dynamics as cells traverse constrictions accurately.

BIOPOINT provides a computational framework for simulating nuclear mechanics and cell-ECM interactions with experimentally derived parameters. By integrating experimental data with a particle-based approach, BIOPOINT offers a practical tool for studying cell behavior that can inform future morphogenetic studies in-vivo or in-vitro.

## 1. Introduction

Morphogenesis is the process by which tissues and organs develop their shape and function, both in vivo during development [1] and in vitro in organoids [2–4] or organs-on-chips [5,6]. This process is controlled by complex biochemical and biomechanical interactions across multiple spatial and temporal scales. Understanding these interactions is crucial for deciphering developmental processes, modeling disease progression, and designing biomimetic engineered tissues. While experimental approaches have provided significant insights, they are often constrained by spatial resolution and temporal control, making it challenging to capture the emergent mechanical behavior of cells and tissues in physiologically relevant contexts [1,7]. One fundamental challenge in morphogenesis is the mechanical interplay between cells, their internal organelles, and their external microenvironment. The nucleus, the largest organelle in the cell, plays a key role in regulating cell migration, differentiation, and mechanotransduction [8,9]. Likewise, the extracellular matrix (ECM) provides biochemical and mechanical cues that guide cellular behavior and emergent tissue structures [10,11]. Therefore, the ability to quantitatively model nuclear mechanics and cell-ECM interactions is critical for understanding how mechanical forces drive morphogenesis.

Computational models provide a powerful means of integrating experimental data and predicting emergent behaviors that are otherwise difficult to capture in vitro [12]. While classical FEM approaches have provided detailed nuclear stress-strain predictions, they typically struggle with large cell deformations and tissue heterogeneity [13,14]. Vertex and Cellular Potts Models (CPM) can simulate migration but have lacked realistic cell and nuclear mechanics [15]. Subcellular Element Modeling (SEM) has emerged as a promising framework for simulating multicellular and subcellular mechanics [16–18]. SEM enables the modeling of emergent shapes by treating cells as ensembles of coarse-grained particles interacting via empirically defined potentials. Recently, we introduced SEM^2^, which allows the calculation of multiscale mechanics in SEM simulations by assessing stress and strain at the particle, cell, and tissue levels [19]. However, a key limitation of existing SEM models is their inability to capture nuclear mechanics and cell-ECM interactions explicitly. Prior SEM implementations have typically represented the nucleus as a single rigid particle, which fails to account for nuclear deformability and the role of intracellular forces in nuclear positioning and chromatin remodeling. Similarly, the ECM has generally been modeled as an implicit solvent rather than a discrete, separate environment directly interacting with cells. Finally, SEM parameters were traditionally derived by homogenizing bulk elastic stiffness and viscosity of non-adherent cells, leading to model parameters that might not be relevant for cell-ECM interactions during morphogenesis. As a result, current SEM frameworks cannot accurately model processes such as nuclear deformation during confined migration or how ECM geometry influences cell spreading and mechanosensing.

To address these limitations, we propose two advancements. First, we introduce a deformable nucleus, represented by phase-separated interacting particles, enabling simulation of nuclear strain and stress distributions. Second, in order to recapitulate *in vitro* experiments, we explicitly represent the ECM as a planar arrangement of static particles interacting with the cell via tunable adhesive potentials. Since everything in the simulation is modeled explicitly via discrete particles, our simulations resemble paintings in the style of famed pointillists Georges Seurat (1859–1891) and Paul Signac (1863–1935). In their honor, we called this framework BIOPOINT. Notably, BIOPOINT enables quantitative predictions of nuclear mechanics and cell-ECM interactions, thanks to the explicit representation of nuclear and ECM particles. Further, the parameters governing potentials can be calibrated directly against experimental data collected in relevant biological conditions, including adherent cells on various ECM environments. To showcase BIOPOINT, we followed a three-step approach. First, we calibrated model parameters using single-cell indentation experiments, ensuring that nuclear and cytoplasmic mechanical properties are quantitatively aligned with experimental force-time data [20,21]. Second, we validated these parameters through cell-spreading simulations on 2D ECM micropatterns, assessing whether BIOPOINT can reproduce experimentally observed relationships between ECM geometry and nuclear shape dynamics [22,23] that were not used for parameter fitting. Finally, we applied the fitted model to the simulations of nuclear deformation during 3D confined migration, testing whether BIOPOINT can replicate experimentally reported nuclear strain and shape changes as cells traverse narrow constrictions [24,25]. This approach aims to demonstrate that BIOPOINT extends to the nucleus and the ECM the ability of SEM to simulate emergent shape and mechanical changes, filling a gap in morphogenesis modeling.

## 2. Results

### BIOPOINT uses heterogeneous discrete particle types to model the cell nucleus and ECM

SEM traditionally treated the nucleus as a single, rigid particle within the cytoplasm, preventing accurate simulation of nuclear deformability and intracellular stress distributions [16,18,19]. Likewise, classical SEM frameworks have implicitly modeled the extracellular matrix (ECM), lacking the ability to capture the influence of ECM geometry on cell spreading and migration. These limitations restrict the ability to model key morphogenetic processes, including nuclear deformation during confined migration and ECM-guided mechanosensing. To overcome these issues, BIOPOINT introduces a deformable nucleus and an explicit ECM representation, as described below.

We first generated a multiparticle nucleus phase separated from the cytoplasmic particles. We initialized a single cell in the simulation as an ensemble of 1000 interacting particles, of which 100 (10%) were designated as nuclear particles (Fig. 1A). These particles were assigned a stronger self-interaction potential (nuclear-nuclear adhesion) than cytoplasmic particles to induce phase separation. The nuclear particles were also assigned a higher stiffness potential (*κ*_*nuc*_= 2× *κ*_0_) to reflect the experimentally observed higher stiffness of the nucleus compared to the surrounding cytoplasm [26] (Fig. 1B). We then performed an equilibration step, allowing the nucleus to coalesce within the cytoplasmic particle ensemble. After equilibration, the nuclear particles successfully segregated into a distinct phase within the cytoplasm, forming a cohesive, deformable nucleus (Fig. 1C). Nuclear and cytoplasmic particles were intermixed in the initial random configuration. Still, after phase separation, fewer than 8% of cytoplasmic particles remained inside the nuclear boundary (Fig. 1C). The nuclear stiffness parameter (*κ*_*nuc*_) effectively maintained the nucleus as a mechanically distinct subregion, consistent with biological observations of nuclear mechanics [26].

**Figure 1.**
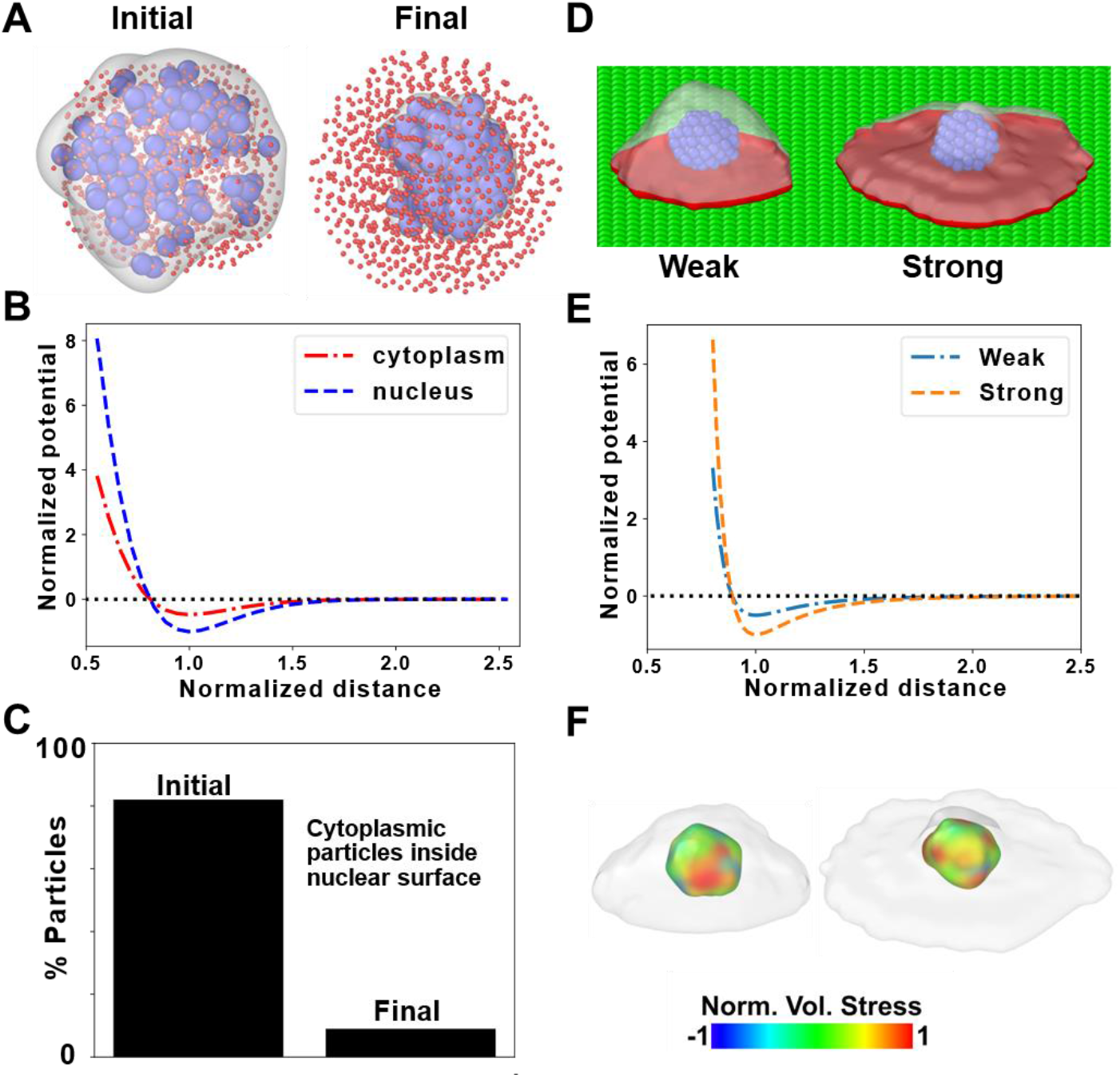
BIOPOINT: Extending SEM^2^ to incorporate a multi-particle nucleus and new particle type for ECM. (A) Representation of the initial and phase-separated structure of an *in silico* cell with Np = 1000, where 900 cytoplasmic (red) and 100 nuclear (blue) particles have been created randomly within a cylindrical space. To visualize the nucleus, we create a surface rendering of nuclear particles. Though the sizes of cytoplasmic and nuclear particles are the same, the former is shown as smaller than the latter for visualization purposes. B) Plot of potential versus distance between cytoplasmic particles and between nuclear particles. C) Percentage of cytoplasmic particles within the nuclear surface in the initial and final samples, with phase-separated nuclear particles. D) Weak and strong interaction with extracellular matrix (ECM) particles can lead to different degrees of cell spreading. Projected surface area at the bottom is shown in red. E) Plot of normalized potential versus distance between ECM and cytoplasmic particles for the weak and strong interactions. F) Spatial distribution of stresses in the nucleus for the weakly and strongly adhering cells, shown in D. The cell membrane is white, and the nucleus is color-coded according to the per-particle stress.

To evaluate the explicit ECM representation, we constructed a ECM layer consisting of static particles arranged in a plane at the base of the simulation domain (Fig. 1D). These ECM particles exerted tunable adhesive interactions with cytoplasmic particles via a Lennard-Jones (LJ) potential. We then systematically varied the cell-ECM adhesion strength to test how this parameter influences cell spreading behavior. Cells on the ECM substrate exhibited distinct spreading behaviors depending on the adhesion potential. When the cell-ECM adhesion strength was low, cells remained compact and retained a near-spherical shape (Fig. 1D, left). Increasing the adhesion strength resulted in greater cell spreading (Fig. 1D, right), with the projected area expanding proportionally to the adhesion potential (Fig. 1E, Fig. S1). Notably, high adhesion levels led to maximal cell spreading, consistent with experimental studies on ECM-guided migration [27,28]. The spatial distribution of stress in the nucleus for the cells with different degrees of spreading can be obtained from per-particle stress computation and visualized (Fig. 1F).

These results confirm that BIOPOINT integrates nuclear deformability and explicit ECM interactions into a SEM framework. Forming a phase-separated nucleus demonstrates that our approach can model intracellular compartmentalization and stress-strain distributions at the nuclear level. Additionally, the planar ECM representation allows for the systematic tuning of adhesion forces, enabling the study of ECM-guided cell behavior. These enhancements overcome key limitations of previous SEM-based models and provide a biophysically relevant platform for studying nuclear mechanics and cell-ECM interactions.

### Calibration of Model Parameters via Single-Cell Indentation Experiments

Previous SEM attempts lacked direct experimental calibration, limiting their accuracy in predicting cellular and nuclear mechanics [16,18,19]. To ensure BIOPOINT accurately captures these properties, we calibrated its parameters using single-cell indentation experiments, a standard technique for measuring cellular viscoelasticity and nuclear stiffness [20,21]. By simulating Atomic Force Microscopy (AFM) indentation and comparing force-time responses with experimental data, we aimed to identify key mechanical parameters using Uncertainty Quantification (UQ) and determine the elastic and viscous properties of the nucleus and cytoplasm using non-linear optimization.

We designed an *in silico* indentation experiment replicating the protocol of Hobson *et al*. [21] (Fig. 2A). A BIOPOINT cell was spread on an ECM layer, with the nucleus positioned near the top surface. To prevent excessive lateral spreading, peripheral cytoplasmic particles were immobilized, mimicking focal adhesions (Fig. 2B). Indentation was performed using a rigid spherical tip moving in a displacement-controlled manner (Supplementary video SV1). In the loading phase, the tip compresses the cell and nucleus. During the hold phase, the tip remains stationary, allowing force relaxation. Finally, in the unloading phase, the tip retracts, relieving compression. Throughout the experiment, force exerted on the tip was recorded, generating force-time curves comparable to AFM data. To simplify the analysis, we assumed no adhesion between the tip and the cell, avoiding artifacts observed in real AFM retraction phases. The spatial distribution of nuclear stress and strain can be computed and visualized during the entire simulation (Supplementary Videos SV2 and SV3).

**Figure 2.**
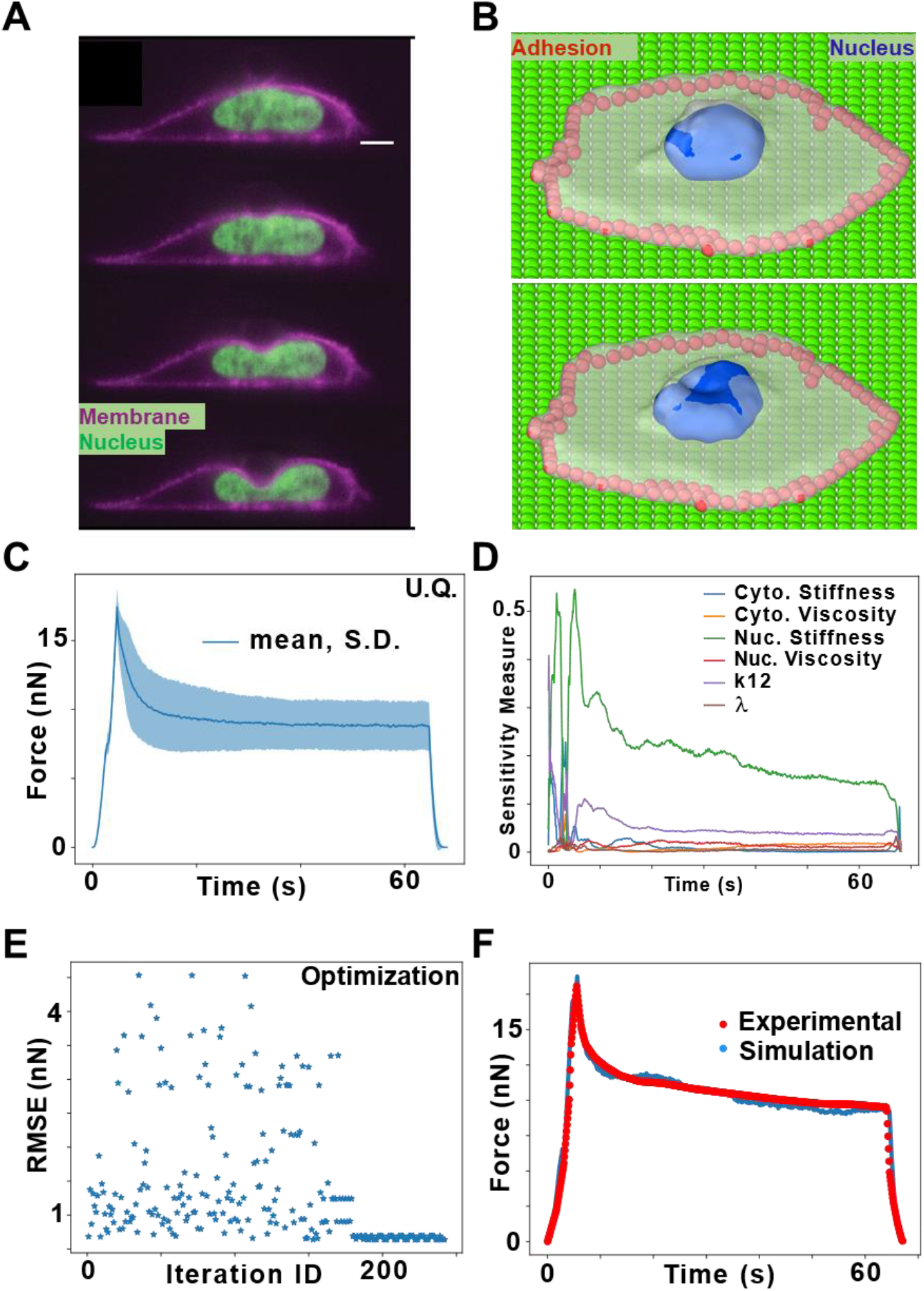
Calibration of model parameters from the spherical indentation of a single cell. A) Fluorescence image of SKOV3 cell undergoing indentation, from Hobson *et al*. [21]. Scale bar: 5 µm. B) Representation of simulated cell, showing cell membrane (white), nucleus (blue), and peripheral constrained particles (red), before and after indentation. C) Uncertainty quantification (UQ) of parameters: Plot of mean and standard deviation of the force values versus time. D) Influence of individual parameters on the force values versus time, as derived from UQ. E) Optimization: Plot of RMSE values for each iteration during the optimization. F) Force-time plot of literature experiment and optimized simulation of single cell indentation.

The simulated force-time curve exhibited a three-phase response, with force increasing during loading, relaxing during the hold phase, and returning to baseline in unloading (Fig. 2C). This response closely matched experimental AFM indentation data [21], confirming that BIOPOINT accurately captures single-cell compression mechanics. Then, we performed UQ to assess which parameters most influence indentation response. Nuclear stiffness and nuclear-cytoplasmic interaction potential (k_12_) had the strongest effect on force-time curves (Fig. 2D). The tuning coefficient (λ) had negligible influence, suggesting that this parameter doesn’t contribute much to cellular and nuclear mechanics. Cytoplasmic viscosity and stiffness contributed but played a secondary role. These results confirm that nuclear mechanics drive indentation responses, reinforcing the need for explicit nuclear modeling in BIOPOINT.

To refine BIOPOINT’s mechanical parameters, we performed non-linear optimization by minimizing the root mean square error (RMSE) between our simulation results and the experimental force-time curve. Upon convergence (Fig. 2E), BIOPOINT learned parameter values that successfully matched experimental force-time curves, confirming that the model’s elastic and viscous properties aligned with real cell behavior (Fig. 2F). Thus, BIOPOINT accurately reproduces experimental force-time curves, identifies dominant mechanical parameters, and optimizes cell mechanics, ensuring a biophysically accurate, experimentally validated framework.

### Validation of BIOPOINT via Cell Spreading on ECM Patterns

Having calibrated BIOPOINT’s parameters, we next validated its ability to predict cell and nuclear shape evolution during ECM-guided spreading. ECM geometry is a key regulator of cell mechanics, influencing morphogenesis, junction remodeling, and spatial organization within tissues [22,29]. We simulated cell spreading on micropatterned ECM substrates to achieve three goals. First, we want to assess whether BIOPOINT reproduces experimentally observed cell shape evolution on different ECM geometries. Second, we want to quantify nuclear shape changes and compare them to known nuclear responses to ECM constraints. Third, we want to isolate the key parameters governing nuclear deformation.

We modeled a single cell spreading on ECM micropatterns with distinct geometries: a circle (Fig. 3A), a square (Fig. 3B), a triangle (Fig. 3C), and a series of elongated rectangles (Fig. 3D, Supplementary video SV4)). The ECM was represented as a layer of immobile particles that interacted with the cytoplasmic particles via adhesive potentials. Surrounding inert particles mimicked non-adhesive substrates (such as the glass on an experimental coverslip), resulting in a confined environment where cells can adhere only to the intended ECM shape. The stress distribution in the spread-out cell can be visualized from the computation of per-particle stress (Fig. S2). Cell spreading was quantified over time using the Cell Shape Index (CSI) and Nuclear Shape Index (NSI), computed as: 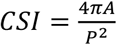 where A is the cell area and P is the perimeter. CSI values near 1 indicate a circular shape, while lower values reflect elongation (Fig. 3E). NSI was measured similarly to track nuclear deformation.

**Figure 3.**
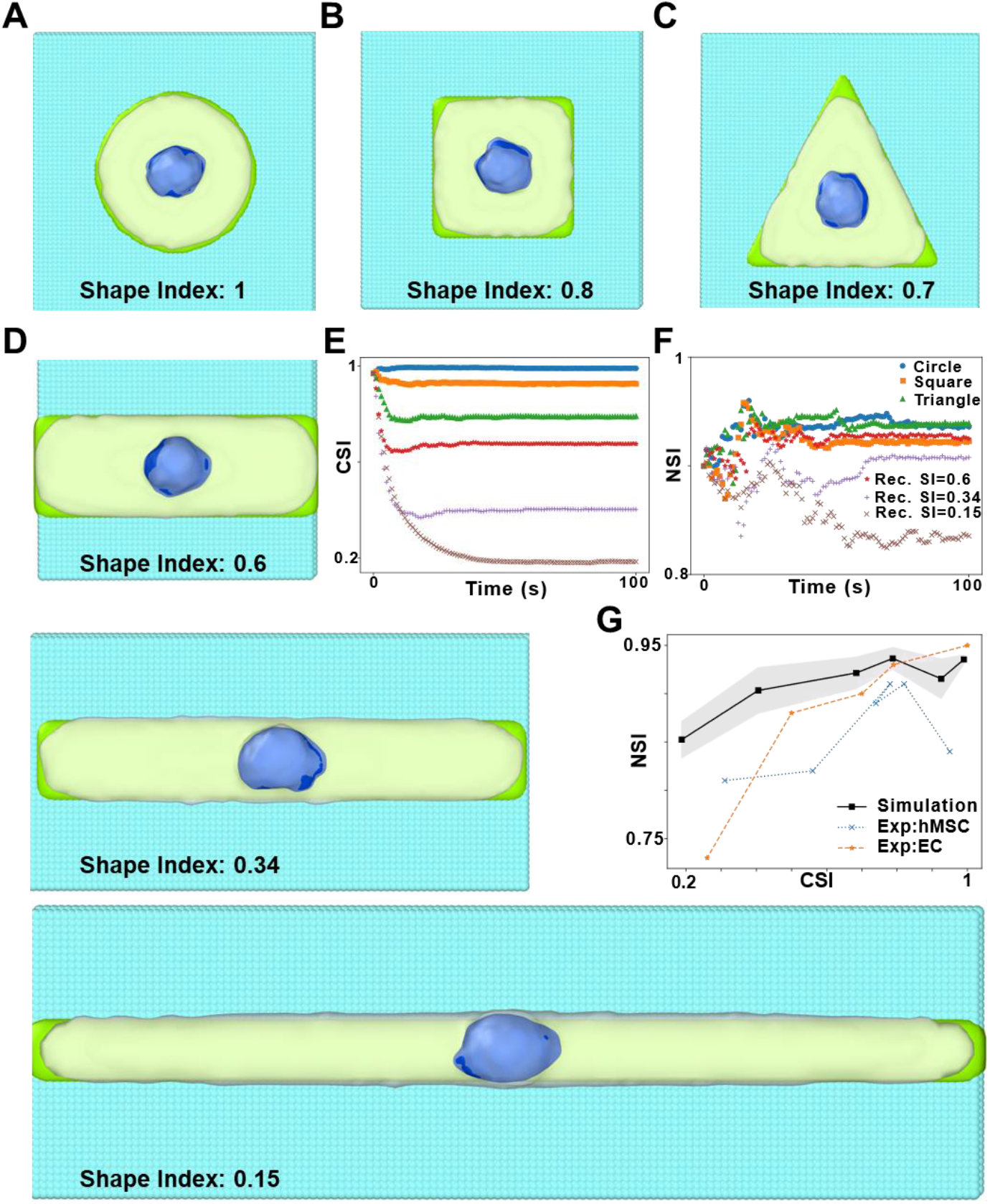
Evolution of shape as cell spreads on various ECM geometries. Simulations of cell spreading on circular (A), square (B), and triangular (C) ECM patterns, where the cell takes up the shape of the pattern. The cell membrane is depicted in white, the nucleus in blue, the ECM surface in green, and surrounding glass particles in cyan. D) Cell spreading on three rectangular patterns with similar area but different shape index. E) Cell shape index (CSI) versus time as the cell spreads on the various ECM patterns. F) Nuclear shape index (NSI) versus time during cell spreading on the various ECM patterns (legends of E and F are the same). G) Plot of NSI versus CSI for our simulations and experimental results from the literature: EC [22] and hMSC [23].

BIOPOINT simulations successfully recapitulated cell spreading behaviors across ECM geometries. Cells spread with maximum circularity on circular ECM, reaching CSI values near 1 (Fig. 3e). Cell circularity reduced in squares and triangles with lower CSI values. The most elongated shape was observed on rectangular ECM, producing minimum CSI values. These trends align with experimental findings that ECM geometry dictates cell morphology [22,23]. While the nucleus in BIOPOINT was deformable, its shape changed minimally in circular, square, and triangular ECM conditions (Fig. 3F). In elongated rectangular ECM, NSI decreased modestly, indicating slight nuclear elongation. The overall NSI trend versus CSI (Fig. 3G) matched experiments, but BIOPOINT underestimated experimentally extreme nuclear deformations. To investigate the discrepancy, we varied the nuclear-cytoplasmic potential strength (k_12_). Increasing k_12_ enhanced nuclear deformability, partially improving experiment agreement (Fig. 3G). However, these simulations failed to capture the drastic nuclear elongation in some experimental conditions, which might require additional potentials to match the stress state in these cell lines (See Fig. S3). Additionally, the stresses in spreading are much smaller than those in indentation, as can be seen from the spatial distribution of stresses and spatial binning plots (Fig. S4), which might non-linearly affect nuclear response.

These findings establish BIOPOINT’s utility for modeling cell-ECM interactions, highlighting future improvements—such as incorporating cytoskeletal constraints—to replicate nuclear deformations better.

### Validation of BIOPOINT in Nuclear Deformation During Constrained Migration

Nuclear deformation is a key factor in cell migration through confined environments, influencing motility, mechanotransduction, and cell fate [11,25]. Experiments have shown that when cells traverse narrow constrictions, their nuclei experience significant shape changes and mechanical stress, which can impact migration efficiency [24,30]. Since BIOPOINT incorporates a deformable nucleus, we tested whether it can reproduce experimentally observed nuclear deformation dynamics during migration through a constriction. By simulating cell movement through a narrow passage flanked by rigid cylindrical constraints, we aimed to validate BIOPOINT’s ability to replicate nuclear shape changes as cells navigate confined spaces. Furthermore, via our analysis of particle-level mechanics, we sought to provide additional mechanical information that would be inaccessible experimentally.

We designed BIOPOINT simulations based on the constricted migration experiments of Keys et al. [24] (Fig. 4A). Two rigid cylindrical obstacles were placed in the simulation domain, forming a narrow constriction. As in our previous work [19], cell migration was driven by addition/removal of cytoplasmic particles at the leading/trailing edge, while nuclear positioning was controlled by a biasing force that maintains nuclear centering within the cell. During the simulation, a cell migrated through the gap, forcing its nucleus to squeeze through the confinement. Using the nuclear shape index (NSI, similar to the CSI discussed above), we tracked nuclear shape over time to show that nuclear deformation matched experimental results (Fig. 4B). We also computed the correlated nuclear ellipticity to show how the nucleus remains rounded (NSI ≈ 1) before entering the constriction (Fig. 4C, t_12_, Supplementary video SV5). However, as the cell moves through the narrow passage (t_2_), the nucleus elongates significantly (NSI decreases, ellipticity increases, Fig. 4D) due to external compression. After exiting (t_3_), the nucleus partially recovers its original shape but retains some deformation, consistent with experimental observations (Fig. 4D, Supplementary video SV5). Next, we leveraged BIOPOINT’s particle-based approach, which allows spatial stress analysis at the nuclear level. As we vary the stiffness of the nucleus, we find higher stress in the stiffer nuclei while passing through the constriction (Fig. 4E, Supplementary video SV6). The absolute values of the stresses are highest in the stiffest nucleus, evident from average stress per particle in spatial bins along the migration direction (Fig. 4F). A similar analysis can be performed for spatial distribution of strain in the cell while passing through the constriction (Fig. S5).

**Figure 4.**
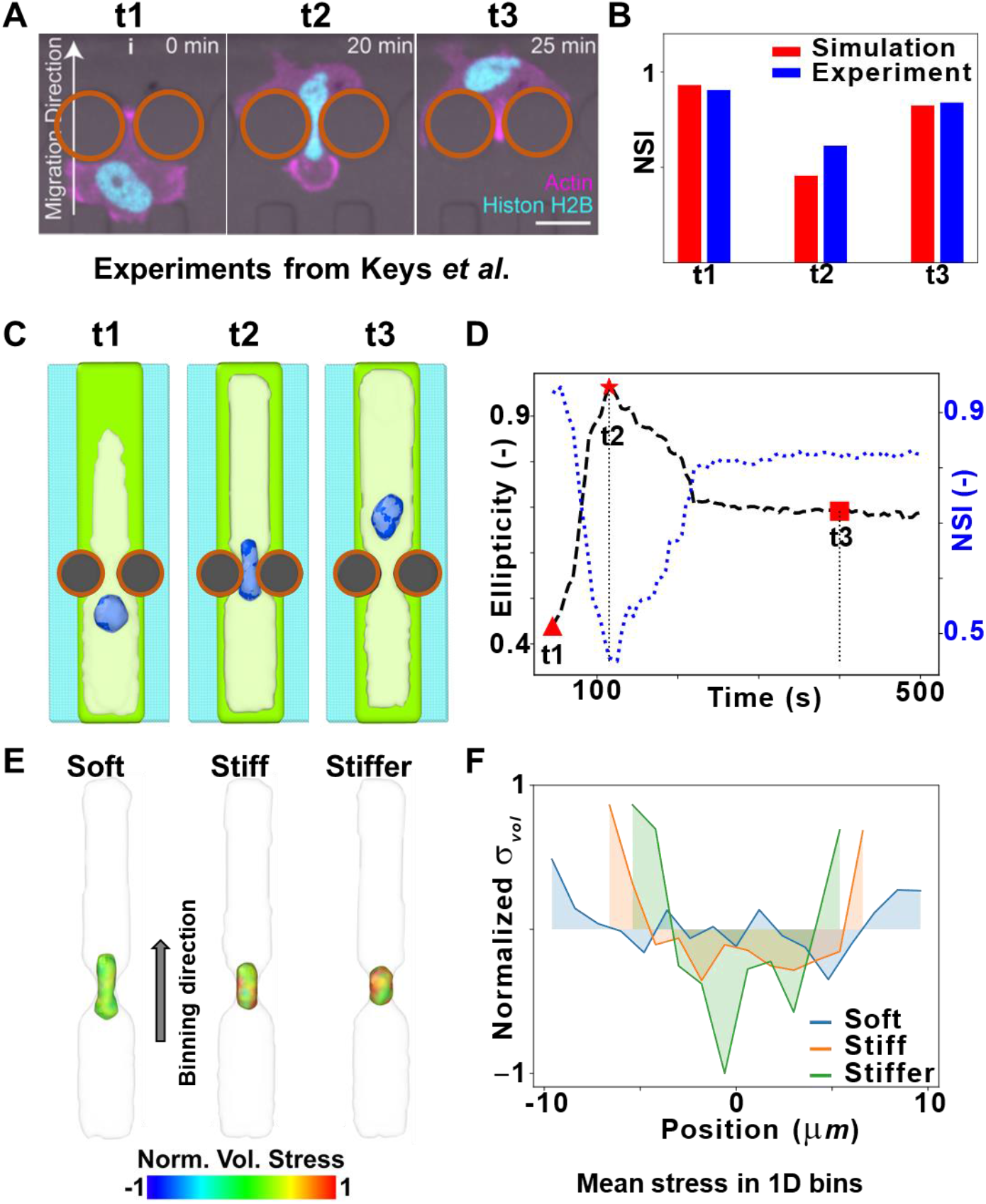
Nuclear deformation when cell passes through a narrow constriction. A) An experimental image sequence of MDA-MB-231 cells migrating through a narrow constriction, as shown by Keys *et al*. [24]. Brown circles are overlaid over the rigid constraints to enhance visualization. Scale bar: 20µm. B) Nuclear shape index (NSI) before (t1), during (t2), and after (t3) passing through the constriction, for our simulations and the experiment shown in (A). C) Simulated images of a nucleus changing shape at various time points as a cell passes through a constriction. As in previous images, the ECM, cell membrane, nucleus, and glass are represented by green, white, blue, and cyan, respectively. Brown circles are overlaid over the rigid constraints to enhance visualization. D) Nuclear ellipticity and NSI versus time as the cell passes through the constriction. E) Visualization of nuclear stress distribution in nuclei with various stiffnesses. The nucleus is color-coded with volumetric stress, and the cell membrane is depicted in white. F) Normalized mean volumetric stress versus position in one-dimensional spatial bins along the direction of migration, indicated by the arrow in E.

These results suggest that nuclear mechanics can contribute to migration efficiency in constrained environments, in good agreement with recent experimental evidence [24]. Additionally, they further validate the ability of BIOPOINT to successfully reproduce nuclear deformation events that spontaneously emerge from cell-ECM interactions in 2D and 3D.

## 3. Discussion

BIOPOINT introduces a coarse-grained, particle-based modeling framework that explicitly represents cytoplasm, nucleus, and extracellular matrix (ECM). By modeling the nucleus as a deformable ensemble of interacting particles (Fig. 1A) and incorporating an explicit ECM representation (Fig. 1D), BIOPOINT enables the study of nuclear mechanics and cell-ECM interactions in ways that were impossible with previous SEM approaches. Our simulations demonstrate that cell shape evolution on ECM micropatterns (Fig. 3), nuclear deformation during confined migration (Fig. 4), and nuclear stress-strain responses under indentation (Fig. 2) align with experimental observations [21–25].

Unlike traditional SEM, BIOPOINT incorporates mechanical parameters calibrated against experiments conducted in realistic scenarios, with cells adhered to isotropic or anisotropic ECM cues. Through single-cell indentation experiments, we determined the elastic and viscous properties of both the nucleus and cytoplasm, allowing precise parameter tuning (Fig. 2). We used Uncertainty Quantification to assess parameter sensitivity, followed by Nelder-Mead optimization, ensuring quantitative agreement between simulated and experimental force-time curves. While uncertainty quantification was computationally efficient, running in parallel on 96-core processors (~420 simulations, ~ 1.5 hours), Nelder-Mead optimization was slower (~2 days, ~240 simulations). Future work could explore parallel computing with Bayesian optimization [31] to improve computational efficiency of optimization.

BIOPOINT’s validation across multiple experimental conditions supports its application for investigating nuclear mechanics in morphogenesis. Our cell spreading simulations confirmed that ECM geometry influences nuclear shape (Fig. 3), though BIOPOINT underestimated nuclear elongation in high-aspect-ratio ECM conditions. This discrepancy likely arises from the absence of actin filament-mediated nuclear compression, which is known to contribute to nuclear deformation [22]. Incorporating polymeric chains to model actin filaments could improve BIOPOINT’s ability to capture cytoskeletal contributions to nuclear reshaping [32,33]. Similarly, simulations of nuclear deformation in confined migration (Fig. 4) successfully reproduced nuclear elongation under compression and partial shape recovery post-constriction, aligning with experimental data [24,25]. However, BIOPOINT does not yet explicitly model cytoskeletal filaments or microtubules, which have been shown to affect nuclear mechanics during migration. Future work could introduce active cytoskeletal components, allowing for a more detailed investigation of nuclear resilience under mechanical stress [32,33]. The number of particles used to model the nucleus could also be varied (thus varying particle size) to investigate its effect on emergent nuclear shape, although preparing a phase separated nucleus could be challenging with increase in the number of particles.

BIOPOINT provides a new tool for studying nuclear mechanics, cell-ECM interactions, and constrained migration, filling a gap between continuum-based finite element models and discrete cell-based approaches such as vertex models and Cellular Potts Models. BIOPOINT extends these classical computational approaches by combining particle-based nuclear mechanics with explicit ECM modeling. Unlike FEM, which excels in stress-strain analysis but struggles with large non-linear deformations [13], BIOPOINT’s particle-based approach allows emergent behavior while retaining mechanical accuracy. Compared to Cellular Potts Models, BIOPOINT explicitly represents nuclear mechanics rather than inferring nuclear deformation indirectly [15]. Finally, while vertex models effectively describe tissue-level behavior, BIOPOINT provides a more detailed subcellular perspective, making it particularly suited for studying nuclear mechanics during migration [7].

Ultimately, we believe that BIOPOINT’s biophysically grounded framework, combined with experimental calibration, makes it particularly useful for biologists and bioengineers investigating emergent nuclear and cell shapes during morphogenesis. While BIOPOINT successfully models nuclear mechanics and cell-ECM interactions, further improvements could enhance its predictive power. Including cytoskeletal constraints would allow more accurate modeling of nuclear flattening under ECM confinement. Additionally, incorporating nuclear envelope rupture events could extend BIOPOINT’s applicability to extreme mechanical stresses observed in confined migration. Finally, future work could explore the integration of biochemical signaling, coupling nuclear mechanics with dynamically regulated stiffness changes.

## 4. Methods

All simulations have been performed utilizing the open-source software, LAMMPS (large-scale atomic/molecular massively parallel simulator), extended by our previously reported open-source package [19]. Similar to our previous work on SEM^2^ [19], visualization is via the licensed version of the software, OVITO [34]. All cells in this work are represented by 1000 particles: 900 cytoplasmic and 100 nuclear particles.

### Preparation of multi-particle nucleus

During sample preparation, initially all 1000 particles were placed randomly in a region of space. The attractive potential between nuclear particles is always stronger (Fig. 1B) than the other potentials. Thus, the nuclear particles attract each other, congregate and eventually form a nucleus separated from the rest of the cell. The strength and cutoff of the nuclear potential is varied during equilibration until the nucleus is separated from the rest of the cell.

### Preparation of ECM layer

The cytoplasm, nucleus and ECM are represented by different particle types. ECM particles were placed on a lattice using the “create_atoms” command, in conjunction with the “lattice” command, in LAMMPS (input scripts provided in GitHub repository). In this simplified representation of ECM, the particles have been made static by setting their velocity to zero and there are no forces acting between them. Thus, they act like a plane of static particles, which attract the cytoplasmic particles of the cell. The attraction between cytoplasmic and ECM particles is modeled by a Lennard-Jones (12-6) potential:

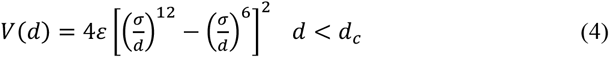

Where σ is the zero-crossing distance, ε is the potential well-depth, and d_c_ is the cutoff distance. The cell spreading can be tuned by adjusting the parameter, ε, of this potential (Fig. 1B).

### Cell spreading

Cell spreading has been carried out by running Brownian dynamics on the cellular particles in the proximity of ECM particles, which results in the cytoplasmic particles moving towards and spreading on the ECM.

### Indentation

For single cell and nuclear indentation, we first performed simulations of cell spreading on an ECM layer with area much larger than the surface area of the cell. The cell height is matched with the experimental setup of Hobson *et al*. [21]. Indentation was implemented by the “fix indent” functionality of LAMMPS with a rigid spherical indenter. During the loading period, the indenter was brought closer to the cell at a fixed rate of 1 µm/s, followed by holding the indenter still and finally retracting the indenter at the same rate. The displacement rate and loading cycle is the same as that used in the experiments. The force exerted on the indenter by each particle is determined by the following relationship:

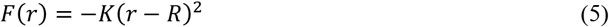

where K is the force constant, r is the distance between the particle and the center of the indenter, and R is the radius of the indenter. The net force on the indenter can be extracted as an output from the LAMMPS software.

### Uncertainty Quantification

In order to determine the impact of individual parameters on the indentation force-time curve, we have performed uncertainty quantification (UQ) and sensitivity analysis [35] using the python library, EasyVVUQ [36,37]. Polynomial chaos expansion with polynomials of order 4 were used in accordance with the recommendations by Eck *et al*. [35] We employed a normal distribution utilizing the package Chaospy [38], with a physically relevant value as the mean (Table 2) and set the standard deviation to 10% of the mean. We performed indentation simulation as described earlier while varying the following parameters: 1. Cytoplasmic elastic stiffness, 2. Cytoplasmic viscosity, 3. Nuclear elastic stiffness, 4. Nuclear viscosity, 5. Tuning coefficient, 6. *k*_*12*_, which is a measure of the relative stiffness of cytoplasmic-nuclear interaction, in comparison to cytoplasmic stiffness.

**Table 1:** Parameters used in the model.

| Parameter | Description | Value |
| --- | --- | --- |
| $R_{cell}$ | Cell radius | 10 $\mu\text{m}$ |
| $m_{cell}$ | Cell mass | 3.1 $\mu\text{g}$ |
| $N_p$ | Total number of particles in a cell | 1000 |
| $N_n$ | Number of nuclear particles | 100 |
| $p_d$ | Sphere close packing density | 0.74 |
| $\rho$ | Scaling factor | 2 |
| $\alpha$ | Shifting factor | 2 |
| $\kappa_0$ | Cell stiffness | $1.2 \times 10^{-2} \text{ N m}^{-1}$ |
| $\eta_0$ | Cell viscosity | $5 \times 10^{-2} \text{ N s m}^{-1}$ |
| $\kappa_{nuc}$ | Nuclear stiffness | $4.5 \times 10^{-2} \text{ N m}^{-1}$ |
| $\eta_{nuc}$ | Nuclear viscosity | $2.1 \times 10^{-1} \text{ N s m}^{-1}$ |
| $\lambda$ | Tuning coefficient | 0.75 |
| $m_p$ | Mass of each particle | 3.1 ng |
| $k_{12}$ | Relative strength of cytoplasmic-nuclear potential | 1.51 |

**Table 2:** Parameter values in UQ and Optimization.

| Parameter | Mean used in UQ | Optimization:<br>Initial Value | Optimization:<br>Final Value |
| --- | --- | --- | --- |
| $\kappa_0$ | $1.5 \times 10^{-2} \text{ N m}^{-1}$ | $1.2 \times 10^{-2} \text{ N m}^{-1}$ | $1.208 \times 10^{-2} \text{ N m}^{-1}$ |
| $\eta_0$ | $5 \times 10^{-2} \text{ N s m}^{-1}$ | $5 \times 10^{-2} \text{ N s m}^{-1}$ | $4.975 \times 10^{-2} \text{ N s m}^{-1}$ |
| $\kappa_{nuc}$ | $6 \times 10^{-2} \text{ N m}^{-1}$ | $4.5 \times 10^{-2} \text{ N m}^{-1}$ | $4.531 \times 10^{-2} \text{ N m}^{-1}$ |
| $\eta_{nuc}$ | $2.2 \times 10^{-1} \text{ N s m}^{-1}$ | $2 \times 10^{-1} \text{ N s m}^{-1}$ | $2.054 \times 10^{-1} \text{ N s m}^{-1}$ |
| $k_{12}$ | 1.5 | 1.5 | 1.51 |
| $\lambda$ | 0.75 | -- | -- |

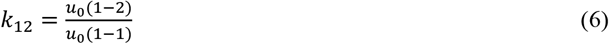

### Optimization

We compute the cost function, which is the root mean squared error (RMSE) between the transient experimental force and the simulated force curve. The parameters which have been optimized are the ones mentioned above (excluding tuning coefficient, λ). The experimental data which has been utilized has been obtained by digitizing the force-time plot of Hobson *et al*. [21] The optimization has been carried out with the Nelder-Mead algorithm from SciPy [39]. The initial values (Table 2) were chosen by analyzing the samples used in the UQ and the impact of individual parameters based on the force curve. In this way, the initial guess of the force curve already matched relatively well with the experimental curve. For convergence, the default tolerance of x values (xatol) and function values (fatol) have been considered, which is 1e-4 for both. The final optimized values of parameters are provided in Table 2. Details of our optimization method and code can be found in our GitHub page (LINK).

### Cell spreading simulations on ECM patterns

For the simulations of cell spreading on ECM patterns, ECM particles were created on a lattice with different planar shapes such as circle, square, triangle and rectangle. Outside the ECM patterns, inert particles have been created to mimic glass substrate of in vitro experiments. The functional form of the potential between glass and cytoplasmic particles is the same as Eq. 4 but the cutoff distance is the zero-crossing distance, i.e. there is only repulsive potential in order to account for volume exclusion. On performing Brownian dynamics on the cellular particles in the proximity of ECM particles, the cellular particles gradually take up the shape of the ECM pattern as shown in Fig. 3.

### Cellular and nuclear shape analysis

The projection of the particle coordinates in a two-dimensional plane of the ECM particles is considered. A convex hull is generated over these 2D coordinates by using the algorithm provided by SciPy. The perimeter (*P*) and surface area (*A*) of the convex hull is then computed. Finally, the cell shape index (CSI) is then determined using:

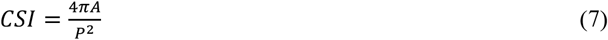

Following a similar procedure, the nuclear shape index (NSI) is computed. For comparison with experiments, we have also plotted values of CSI versus NSI of a couple of cell lines.

### Cell migration through constriction

For the simulations involving a cell passing through cylindrical obstructions, we carried out migration via the addition and removal of particles on a rectangular ECM pattern (Input script available on GitHub). A biasing force on the nucleus towards the cell center on a two-dimensional plane repositions it throughout the migration simulation [19]. The cylindrical obstructions have been implemented by rigid indenters with “fix indent” functionality of LAMMPS.

The NSI is computed by following the procedure mentioned above (similar to CSI). The NSI of the experimental images is measured by digitizing the shape of the nucleus and generating a convex hull over the coordinates.

### Nuclear ellipticity computation

For nuclear shape analysis, a three-dimensional ellipse has been fitted over the coordinates of nuclear particles using an existing Python script [40]. The ellipticity of the fitted ellipse, *f*, is measured using the relation:

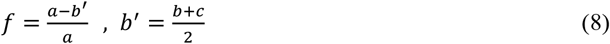

Where *a* is the largest radius of the fitted ellipse and *b* and *c* are the two smaller radii.

### Visualization, stress and strain computations

Generating a surface mesh over the particles has been carried out in OVITO [34], employing a Gaussian density method. The per-particle stress can be computed utilizing the “compute stress/atom” functionality of LAMMPS. By transferring the particle-level stress on the enclosing surface in OVITO, we could visualize the stress distribution in the nuclear surface mesh. The average volumetric stress per particle in 1D spatial bins then provided us with a measure of the nuclear stress distribution along the direction of binning. Additionally, the per-particle shear strain of the nuclear particles with respect to the initial configuration was computed via the “Atomic Strain” modifier of OVITO [34].

## Supporting information

BIOPOINT Supplementary Information

Single cell indentation simulation

Nuclear stress during indentation

Nuclear strain during indentation

Cell spreading on patterns

Cell migration through constriction

Nuclear stress while passing through constriction

## Supplementary material

- **Supplementary figures:**

º Figure S1. Quantification of spreading by measuring projected surface area of cells.
º Figure S2. Spatial stress distribution in the bottom surfaces of cells on patterns.
º Figure S3: Drastic nuclear shape change in rectangular patterns.
º Figure S4: Distribution of nuclear stress during indentation and spreading simulations.
º Figure S5: Spatial strain analyses for cell migration through a constriction.
- **Supplementary videos:**

º Video S1: AFM indentation of a cell with nucleus.
º Video S2: Nuclear stress during indentation.
º Video S3: Nuclear strain during indentation.
º Video S4: Cell spreading on patterns.
º Video S5: Cell migrating through a constriction.
º Video S6: Nuclear stress while passing through a constriction.

## Acknowledgments

This work was supported by the European Research Council (ERC) Starting Grant #852560 to FSP. We would like to thank Constanze Kalcher and Alexander Stukowski of OVITO GmbH for the discussions which benefitted several of our post-processing tasks, including quantifying the number of cytoplasmic particles inside the surface mesh encompassing nuclear particles.

## Data availability statement

The data that supports the findings of this study are available on Zenodo (Link for reviewers, please note that we will finalize the repository during / after the review process to share the latest data) and the code is maintained on GitHub: https://github.com/Synthetic-Physiology-Lab/biopoint

## Ethics approval statement

Not required.

