## Supplementary material for "BIOPOINT: A particle-based model for probing nuclear mechanics and cell-ECM interactions via experimentally derived parameters": BIOPOINT Supplementary Information

### ABSTRACT

Morphogenesis involves biochemical and biomechanical interactions across multiple spatial and temporal scales. Experimental studies alone struggle to resolve these dynamics, necessitating computational models. Among these, subcellular element modeling (SEM) has proven helpful in simulating cellular and tissue-scale emergent behaviors. However, traditional SEM frameworks lack explicit representations of nuclear mechanics and cell-extracellular matrix (ECM) interactions, limiting their ability to capture key biology.

Here, we introduce **BIOPOINT**, a particle-based computational framework that extends SEM by incorporating (1) a deformable, multi-particle nucleus to simulate nuclear stress and strain distributions and (2) an explicit ECM representation using a structured array of static particles interacting via tunable adhesive potentials. To ensure biological relevance, we calibrated BIOPOINT's parameters against single-cell indentation experiments, overcoming prior limitations of ad hoc parameter selection in SEM. We validate BIOPOINT by comparing simulated cell behaviors to experimental observations in three key scenarios: (i) single-cell indentation, demonstrating agreement with force-time curves from atomic force microscopy (AFM) studies; (ii) cell spreading on ECM micropatterns, confirming that nuclear deformation follows ECM constraints; and (iii) nuclear deformation during confined migration, showing BIOPOINT predicts nuclear shape dynamics as cells traverse constrictions accurately.

BIOPOINT provides a computational framework for simulating nuclear mechanics and cell-ECM interactions with experimentally derived parameters. By integrating experimental data with a particle-based approach, BIOPOINT offers a practical tool for studying cell behavior that can inform future morphogenetic studies in-vivo or in-vitro.

#### SUPPLEMENTARY INFORMATION

##### 1. Quantification of spreading by measuring projected surface area of cells

The degree of cell spreading on an ECM substrate depends on the strength of interaction between the cell and ECM. (Fig. S1A). This can be quantified by measuring the projected surface area of the cell (Fig. S1B). The steady state projected area after cell spreading is much higher for the strongly interacting cell-ECM.

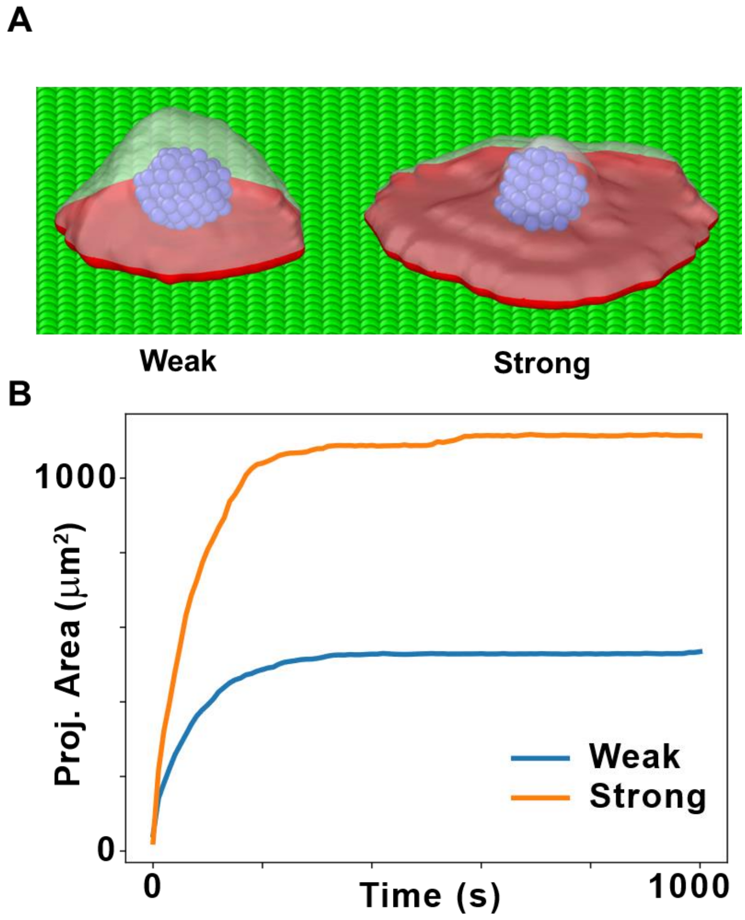


**Figure S1. Projected surface area of cells spreading on ECM.** (A) Weak and strong interaction with extracellular matrix (ECM) particles can lead to different degrees of cell spreading. The projected surface area is indicated in red. B) Plot of the projected surface area of the cell as it spreads on ECM, versus time.

##### 2. Spatial stress distribution in the bottom surfaces of cells on patterns

In the cell spreading simulations on circular (Fig. S2A), square (Fig. S2B) and triangular (Fig. S2C) ECM patterns, mean volumetric stress per particle was computed utilizing LAMMPS "compute stress/atom" functionality. The particles and enclosing surfaces have been color coded as per the normalized mean volumetric stress. One can see on the bottom surface of the stretched-out cells that the peripheral region is under tensile (positive) stress whereas the inner region is under compressive (negative) stress.

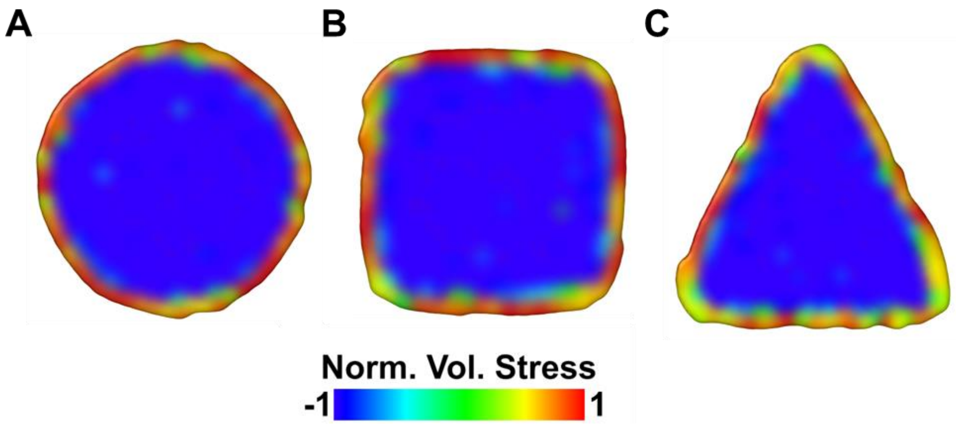


**Figure S2. Spatial stress distribution in the bottom surfaces of cells on patterns.** Visualization of spatial distribution of normalized mean volumetric stress along the bottom surface of cells, which are spread on A) circular, B) square and C) triangular ECM patterns.

##### 3. Drastic nuclear shape change in rectangular patterns

The experimental observation of drastic nuclear shape change in certain cell types such as the U2OS could not be captured with our calibrated parameters obtained from the indentation simulations (Fig. S3A). This could be because of the lack of an explicit treatment of actin filaments in our model. However, on including a potential between the nucleus and ECM particles (2-3 potential), we observe that we could simulate scenarios of drastic shape change of the nucleus (Fig. S3B). Though there is no direct connection between the ECM and the nucleus in reality, this might indicate an indirect connection between the nucleus and ECM in certain cell types where drastic nuclear shape change is observed during spreading. The cumulative effect of the actin filaments in these cell types might be captured through a 2-3 potential in our model.

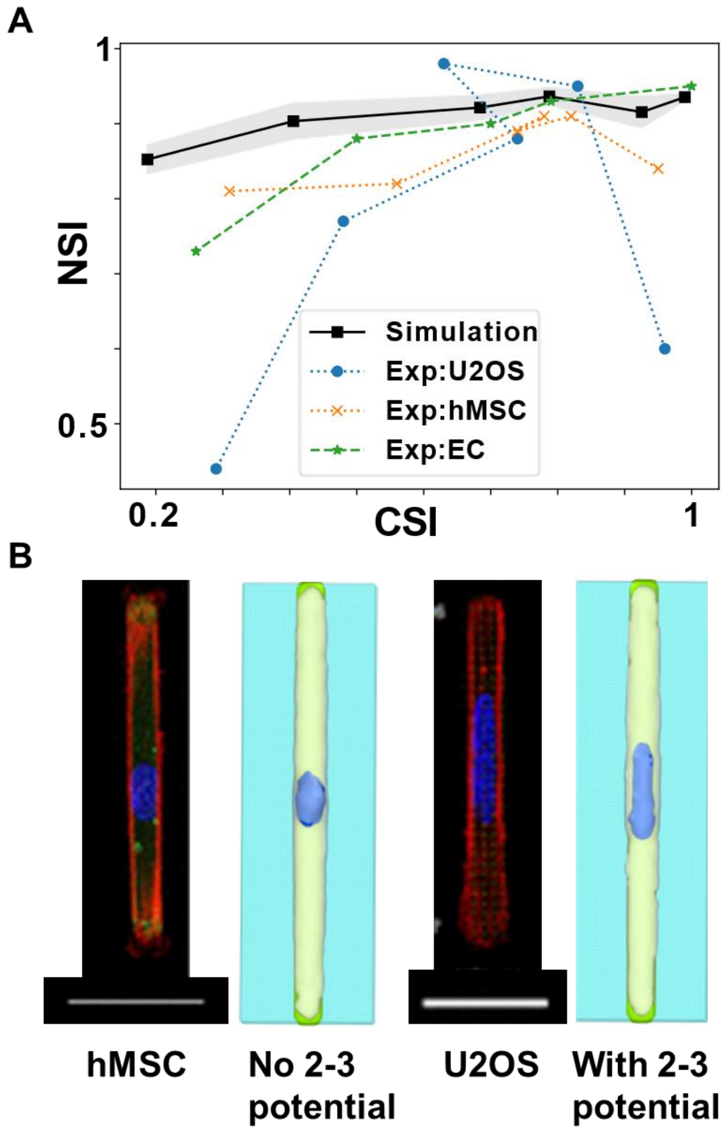


**Figure S3. Effect of the nuclear-ECM potential.** A) Plot of NSI versus CSI for representative simulations and experimental results from the literature for cell types: EC [22], hMSC and U2OS [23]. B) Comparison of cell spreading simulations versus experimental results for hMSC and U2OS cells [23]. Experimental representative widefield images from Sarikhani *et al*. [23] have ECM in green, actin in red and nucleus in blue. Scale bar is 50 µm and 40 µm for hMSC and U2OS cells respectively. The simulations of cell spreading are without and with nucleus-ECM (2-3) potential, respectively

#### 4. Distribution of nuclear stress during indentation and spreading simulations

One of the advantages of BIOPOINT is that the stress distribution can be visualized at all times from per-particle stress computation. The visualization of nuclear stress in the spreading (Fig. S4A) and indentation (Fig. S4B) simulations show that there are considerably higher stresses in the nucleus during indentation. This can be further corroborated from the plot of average stress per particle in spatial bins along the x axis (Fig. S4C). The absolute values of stresses are much more in the indented nucleus as compared to the nucleus in the spread-out cell.

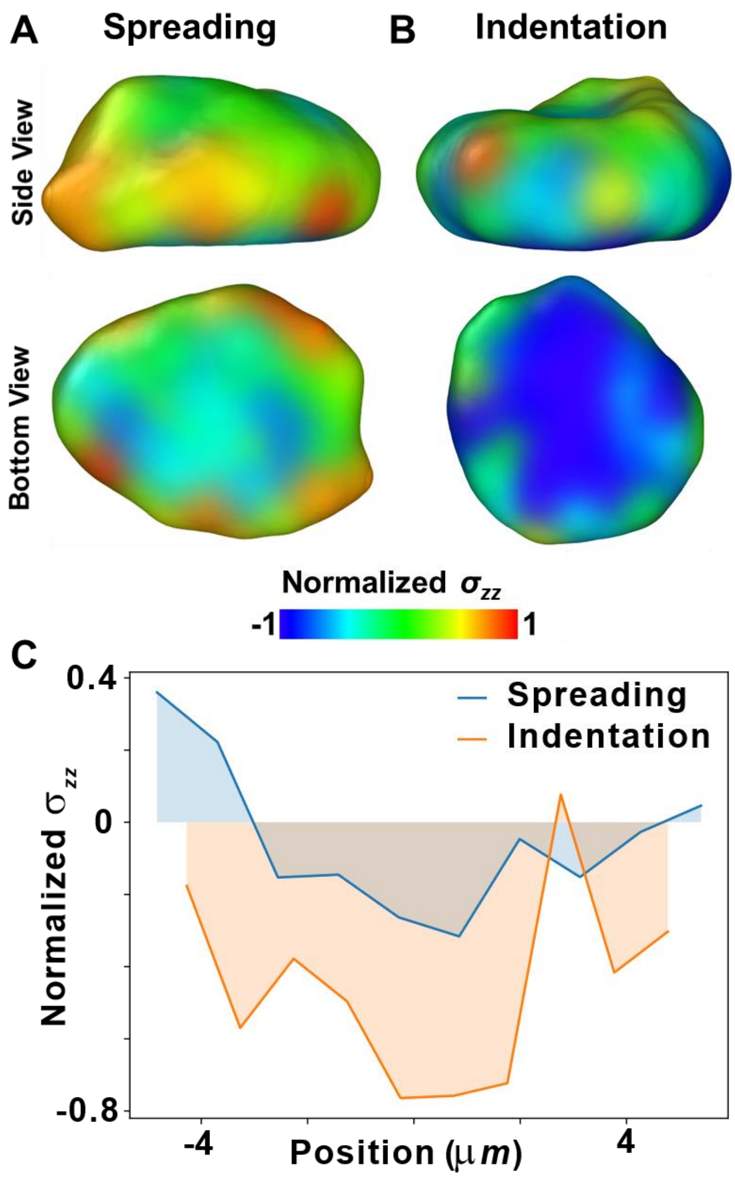


**Figure S4. Distribution of nuclear stress during indentation and spreading simulations.** Side view and the bottom surface view of A) a nucleus of a cell spread on ECM substrate and B) an indented nucleus. The nucleus has been color coded with normal component of the per-particle stress tensor ($\sigma_{zz}$). C) Plot of normalized $\sigma_{zz}$ versus position along one-dimensional spatial bins along the x axis.

##### 5. Spatial strain analyses for cell migration through a constriction

As the cell moves through the narrow passage (Fig. S5A, $t_2$), the nucleus elongates significantly (NSI decreases, ellipticity increases) due to external compression (Fig. S5B). After exiting ($t_3$), the nucleus partially recovers its original shape but retains some deformation (Fig. S5B), consistent with experimental observations. We leveraged BIOPOINT's particle-based approach, which allows spatial strain analysis even at the nuclear level. We computed per-particle shear and volumetric strain, revealing that the maximum shear strain occurs while the nucleus is inside the constriction (Fig. S5C). Further, a residual strain persists even after passage, indicating incomplete shape recovery (Fig. S5D). Finally, a strong correlation exists between nuclear ellipticity and average strain, confirming that nuclear deformation is mechanically driven (Figs. S5B, D).

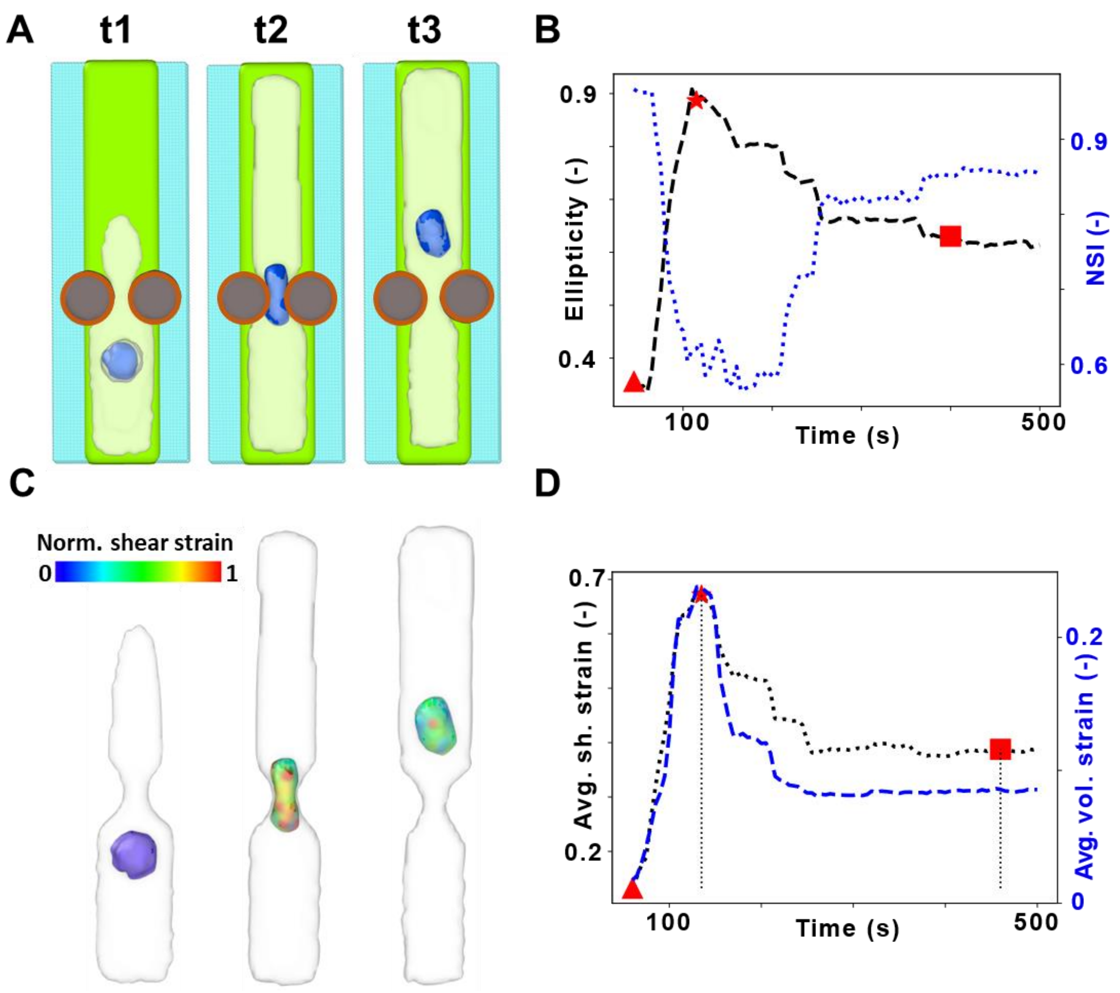


**Figure S5. Nuclear strain when cell passes through a narrow constriction.** A) Simulated images of a nucleus changing shape at various time points, as a cell passes through a constriction. As in previous images, the ECM, cell membrane, nucleus and glass are represented by the colors green, white, blue and cyan, respectively. Brown circles are overlaid over the rigid constraints to enhance visualization. B) Nuclear ellipticity and NSI versus time as the cell passes through the constriction. C) Nucleus color-coded with shear strain shows how it is strained as it passes through the constriction. Cell membrane is depicted in white. D) The mean shear and volumetric strain per nuclear particle go through a maximum as the nucleus passes through the constriction.
